# A comparative study of machine learning algorithms in predicting severe complications after bariatric surgery

**DOI:** 10.1101/376038

**Authors:** Yang Cao, Xin Fang, Johan Ottosson, Erik Näslund, Erik Stenberg

## Abstract

Accurate models to predict severe postoperative complications could be of value in the preoperative assessment of potential candidates for bariatric surgery. Traditional statistical methods have so far failed to produce high accuracy. To find a useful algorithm to predict the risk for severe complication after bariatric surgery, we trained and compared 29 supervised machine learning (ML) algorithms using information from 37,811 patients operated with a bariatric surgical procedure between 2010 and 2014 in Sweden. The algorithms were then tested on 6,250 patients operated in 2015. Most ML algorithms showed high accuracy (>90%) and specificity (>0.9) in both the training and test data. However, none achieved an acceptable sensitivity in the test data. ML methods may improve accuracy of prediction but we did not yet identify one with a high enough sensitivity that can be used in clinical praxis in bariatric surgery. Further investigation on deeper neural network algorithms is needed.

## Introduction

Morbid obesity is a global public health threat of growing proportions(Ng et al., 2014). Bariatric surgery offers the best chance for long-term weight-loss and resolution of comorbidities(Sjostrom et al., 2004). Although modern bariatric surgery is considered to be safe, severe postoperative complications still occur(Finks et al., 2011; Stenberg et al., 2014). Accurate prediction models for severe postoperative complications could aid preoperative decision making for surgeons, anesthesiologists and patients. These models could also serve as basis for case-mix comparisons between different centers. Some prediction models based on linear regression of patient-specific data allow for relatively simple and interpretable inference; however, they have so far been proven inaccurate and can thus not be used in clinical practice(Geubbels et al., 2015; Stenberg et al., 2018).

In contrast, some machine learning (ML) methods have been shown to provide quite accurate predictions, and have increasingly been used in diagnosis and prognosis of different diseases and health conditions(Anderin et al., 2015; Kourou et al., 2015; Pan et al., 2017). ML methods are data-driven analytic approaches that specialize in the integration of multiple risk factors into a predictive algorithm(Passos et al., 2016). Over the past several decades, ML tools have become more and more popular for medical researchers. A variety of ML algorithms, including artificial neural networks, decision trees, Bayesian networks, and support vector machines (SVMs) have been widely applied with the aims to detect key features of the patient conditions and to model the disease progression after treatment from complex health information and medical datasets. The application of different ML methods for feature selection and classification in multidimensional heterogeneous data can provide promising tools for inference in medical practices. These highly nonlinear approaches have been utilized in medical research for the development of predictive models, resulting in effective and accurate decision making(Ali, 2017; Jiang et al., 2017).

Although new and improved software packages have significantly eased the implementation burden for many ML methods in recent years, few studies have used ML methods to examine the risk factors or predict the prognosis after bariatric surgery, including diabetes remission(Hayes et al., 2011; Pedersen et al., 2016), complication(Razzaghi et al., 2017), weight status(Piaggi et al., 2010; Thomas et al., 2017), and adverse events and death(Ehlers et al., 2017). Even though there is evidence that the use of ML methods can improve our understanding of postoperative progression of bariatric surgery, an appropriate level of validation is needed in order for these methods to be considered in the clinical practice.

In this study, we compared different conventional supervised ML algorithms in the modeling of severe postoperative complication after bariatric surgery. The study was based on the data from the Scandinavian Obesity Surgery Registry (SOReg). The SOReg is a national quality and research register, covering virtually all bariatric surgical procedures performed in Sweden since 2010. The register has been described in detail elsewhere(Hedenbro et al., 2015; Stenberg et al., 2014), and a prediction model based on logistic regression for the same group of patients has been described previously(Stenberg et al., 2018). The aim of the current study was to find an algorithm or algorithms that perform well not only on the training data but also on the test data that were not used to train the algorithms.

## Results

Baseline characteristics of the patients in the training data and the test data are presented in Tables 1 and 2. The percentages of severe complication in the two data sets are 3.2% and 3.0%, respectively. No statistically significant difference was found for percentages of severe complication between the two data sets (Pearson chi-square = 0.8283, p = 0.363).

Univariable analyses indicate that differences of mean age, mean body mass index (BMI), median HbA1c, percentages of comorbidities for hypertension, diabetes, dyslipidaemia, and previous venous thromboembolism, and percentage of revisional surgery between the patients presenting and without severe complication are statistically significant in the training data (Table 1). In the test data, the statistically significant differences were found for age, waist circumference (WC), HbA1c, dyslipidaemia, and revisional surgery (Table 2).

Multivariable logistic regression analysis for the same data was published elsewhere(Stenberg et al., 2018). In brief, revisional surgery, age, low BMI, operation year, WC, and dyspepsia were associated with the an increased risk for severe postoperative complication, however, the performance of the multivariable logistic regression model for predicting the risk in individual patient case was poor. Validation of the model tested on patients operated in 2015 resulted in an area under the receiver operating characteristic (ROC) curve of only 0.53, a Hosmer-Lemshow goodness of fit 17.91 (p=0.056) and Nagelkerke R^2^ 0.013(Stenberg et al., 2018).

In current study, 19 supervised machine learning algorithms were compared and ten of them were also trained using the synthetic minority oversampling technique (SMOTE), resulting in 29 ML algorithms. Most of the machine learning algorithms shown high accuracy (>90%) and specificity (>0.9) for both training data and test data (Table 3), except that bagging linear discriminant analysis (LDA), bagging quadratic discriminant analysis (QDA), adaptive boosting (AdaBoost) support vector machine (SVM), and multilayer perceptron (MLP) shown low accuracy (<60%) for SMOTE training data, and oversampling-based bagging QDA shown low accuracy for test data (accuracy = 56.1%) (Table 3).

Although most of the algorithms shown low sensitivity for both the training data and the test data, some of them exhibited promising prediction ability in the training data. Sensitivities of oversampling-based bagging QDA, random forest, AdaBoost extremely randomized (AdaExtra) trees, AdaBoost gradient regression (AdaGradient) trees, bagging k-nearest neighbor (KNN), and deep learning neural network (NN) are 0.707, 0.965, 0.980, 0.968, 0.996, and 0.757 for SMOTE training data, respectively (Table 3). Even for test data, oversampling-based bagging QDA and AdaBoost SVM show significant higher prediction ability than other algorithms. The sensitivities of the two algorithms are 0.417 and 0.364, respectively (Table 3). However, they still do not achieve an acceptable level for practical application.

When considering sensitivity and specificity together, most of the algorithms did not show better prediction ability than a random predictor, i.e. an area under ROC curve of 0.5. The areas under the ROC curves for all the algorithms, except for oversampling-based random forest, AdaExtra trees, and adaGradient trees, and KNN, are around 0.5 (Figures 1 - 4). Although oversampling-based random forest, AdaExtra trees, AdaGradient trees, and KNN show outstanding prediction ability on the SMOTE training data (areas under ROC curves are above 0.9), their performance on the test data are not optimistic (Figures 2 and 3).

The performance of the three regression-based algorithms (logistic regression, LDA, QDA), SVM, and the two neural network-based algorithms (MLP and deep learning NN) was poor in any situation. However the bagging MLP and deep learning NN outperforms the tree-based algorithms (Figures 2 and 4) for test data, their areas under ROC curves for the test data are 0.58 and 0.56, respectively (Figure 4) that are greatest among all the algorithms.

## Discussion

Historically, laparoscopic gastric bypass has for a long time been the most common bariatric procedure in Sweden, although laparoscopic sleeve gastrectomy has increased in popularity over more recent years(Stenberg et al., 2014; The international federation for the surgery of obesity and metabolic disorders, 2017). The surgical technique is highly standardized with more than 99% of all gastric bypass procedure being the antecolic, antegastric, laparoscopic gastric bypass (so called Lönnroth technique)(Olbers et al., 2003). Virtually all patients receive pharmacologic prophylaxis for deep venous thrombosis and intraoperative antibiotic prophylaxis(Hedenbro et al., 2015; Stenberg et al., 2014). Patients who have bariatric surgery are exposed to the risk of having postoperative complications, which may increase the complexity of managing safety and healthcare costs.

Previous studies on postoperative complications of bariatric surgery have mainly used scoring for identifying patients who are more likely to have complications after surgery. However, these methods are not sensitive enough for clinical application(Geubbels et al., 2015; Stenberg et al., 2018). The potential of ML tools as clinical decision support in identifying risk factors and predicting health outcomes is therefore worth investigation on complications associated with bariatric surgery. To our knowledge, there is only one study that compared the performance of different ML algorithms in predicting the postoperative complications in imbalanced bariatric surgery data set(Razzaghi et al., 2017). Although the study indicates that the combination of a suitable feature selection method with ensemble ML algorithm equipped with SMOTE can achieve higher performance in predictive models for bariatric surgery risks, the ML algorithms were not validated using external test data. After all, for prediction purpose, we are not very interested in whether or not an algorithm accurately predicts severe complication for patients used to train the algorithm, since we already know which of those patients have severe complications, but are interested in whether the algorithms may accurately predict the future patients based on their clinical measurements.

Our study compared in total 29 ML algorithms using real world data. Although the sensitivities of the algorithms were generally low, the study indicates that some ML algorithms were able to achieve higher accuracy than tradition logistic regression models(Geubbels et al., 2015; Stenberg et al., 2018). Four of 29 algorithms were able to achieve high sensitivity (>0.95) and two achieved moderate sensitivity (>0.70) in the training data, including three tree-based algorithms, bagging KNN, bagging QDA, and deep learning NN. We should notice that all the high or moderate sensitivities were obtained from SMOTE training data and/or using ensemble algorithms. Our findings support the previous study that ensemble ML algorithms equipped with SMOTE can achieve higher performance metrics for imbalanced data(Razzaghi et al., 2017).

Despite showing promising capability of prediction in training data, none of the 29 ML algorithms satisfactorily predicted severe postoperative complication after bariatric surgery in the test data. Why did the algorithms do a poor job of predicting the patients who had severe complication in test data? One potential explanation for this may be related to the limited number of severe postoperative complication in the current dataset, which cannot reveal the underlying relationship between risk factors and adverse health outcomes. Although there are several known risk factors, each of them only imposes a small increase in the risk for postoperative complication(Finks et al., 2011; Longitudinal Assessment of Bariatric Surgery et al., 2009; Maciejewski et al., 2012; Stenberg et al., 2014). Another likely explanation may be that preoperatively known variables are insufficient to predict postoperative complications. In previous studies, the highest accuracy for prediction of postoperative complication has been models including operation data, mainly intraoperative complication and conversion to open surgery(Geubbels et al., 2015; Stenberg et al., 2014). Although including intraoperative adverse events and conversion to open surgery may improve the accuracy of prediction models, such models would not be useful in the preoperative assessment for patients or for case mix comparisons. Furthermore, because the algorithms try to minimize the total error rate out of all classes, irrespective of which class the errors come from, they are not appropriate for imbalanced data such as what we used in our study(Maalouf et al., 2018).

Compared with traditional generalized linear predictive models, non-linear ML algorithms are more flexible and may achieve higher accuracy but at the expense of less interpretability. Although there are interpretable models such as regression, Naïve Bayes, decision tree and random forests, several models are not designed to be interpretable(James et al., 2013). The aim of the methods is to extract information from the trained model to justify their prediction outcome, without knowing how the model works in details. The trade-off between prediction accuracy and model interpretability is always an issue when we have to consider in building a ML algorithm. A common quote on model interpretability is that with an increase in model complexity, model interpretability goes down at least as fast. Fully nonlinear methods such as bagging, boosting and support vector machines with nonlinear kernels are highly flexible approaches that are harder to interpret. Deep learning algorithms are notorious for their un-interpretability due to the sheer number of parameters and the complex approach to extracting and combining features. Feature importance is a basic (and often free) approach to interpreting the model. Although some nonlinear algorithms such as tree-based algorithms (e.g. random forest) may allow to obtain information on the feature importance, we cannot obtain such information from many ML algorithms.

Therefore, recent attempts have been made to improve interpretability for the black-box algorithms even such as deep learning. Local interpretable model-agnostic explanations (LIME) is one of them to make these complex models at least partly understandable. LIME is a more general framework that aims to make the predictions of ‘any’ ML model more interpretable. In order to remain model-independent, LIME works by modifying the input to the model locally(Mishra et al., 2017; Ribeiro et al., 2016). So instead of trying to understand the entire model at the same time, a specific input instance is modified and the impact on the predictions are monitored.

Regarding specific algorithm, though their motivations differ, the logistic regression and LDA or QDA methods are closely connected, therefore we were not surprised that LDA or QDA did not show significant improvement in prediction than logistic regression(Stenberg et al., 2018). KNN takes a complete different approach from classification which is completely non-parametric(James et al., 2013). Therefore, we can expect it to outperform parametric models such as logistic and LDA. However, KNN cannot tell us which predictor are of importance. QDA serves as a compromise between the non-parametric KNN and the LDA and logistic regression. Though not as flexible as KNN, QDA can perform better in the limited training data situation. MLP is a class of feedforward artificial neural network, which consists of at least three layers of nodes. Its multiple layers and non-linear activation can distinguish data that is not linearly separable. Deep learning NNs are high-level NNs including convolutional NN and recurrent NN et al. In our study, the deep learning NN with five hidden layers outperforms the conventional MLP with two hidden layers, especially on SMOTE training data (areas under ROC curves are 0.67 vs. 0.37), which deserves further investigation in the future.

Our study demonstrates that ensemble learning may improve predictions by combining several base algorithms. However, usually there are several ensemble methods available, such as bagging, boosting, and stacking(Zhou, 2012). A number of studies have shown that, when decomposing a classifier’s error into bias and variance terms, AdaBoost is more effective at reducing bias, bagging is more effective at reducing variance, and stacking may improve predictions in general(Kotsiantis et al., 2007). There is no golden rule on which method works best. The choice of specific ensemble methods is case by case and depends enormously on the data.

There are some limitations in our study. First, the study was limited to data registered within the SOReg. Cardiovascular and pulmonary comorbidities other than sleep apnea are not mandatory variables within the registry and could thus not be included in the model. Although these comorbidities are known risk factors for postoperative complications(Finks et al., 2011; Gupta et al., 2011; Maciejewski et al., 2012), they are not highly prevalent in European studies(Geubbels et al., 2015). Second, although we compared 29 ML algorithms investigated in our study, they are convenient and feasible methods for general medical researchers. Because of computational complexity and less interpretability, many complicated and advanced ML algorithms were not yet investigated in our study. However, our study at least points out a promising way for future investigations, i.e. deep learning NN equipped with SMOTE. Last but not the least, the exhaustive grid search was used in our hyperparameter optimization, which is extremely resource consuming and not optimal for complex ML algorithms, therefore other advanced methods such as gradient-based or evolutionary optimization would be considered in the future.

## Conclusion

ML algorithms have the potential to improve the accuracy in predicting the severe postoperative complication among the 44,061 Swedish bariatric surgery patients during 2010 - 2015. Because the imbalance nature of the data where the number of the interested outcome is relative small, oversampling technique needs to be adopted to balance the two outcomes (presenting or without severe complication). Ensemble algorithms outperform base algorithms. In general, deep learning NN results in better predictions for unseen patients.

## Materials and Methods

### Patients and features

Patients registered in the SOReg between 2010 and 2015 were included in the present study. All patients who underwent a bariatric procedure between 2010 and 2014 were used as training data in the ML. Data from patients who underwent a bariatric surgical procedure in 2015 were used as test data to validate the algorithm’s performance in predicting sever postoperative complication within 30 days after surgery. In total, 37,811 and 6,250 bariatric patients from SOReg were included in the training data and test data, respectively. In total 16 features were included in ML, including five continuous features (age, HbA1c, body mass index [BMI], waist circumstance [WC]), and operation year) and 11 binary features (sleep apnoea, hypertension, diabetes, dyslipdaemia, dyspepsia, depression, musculoskeletal pain, previous venous thromboembolism, revisional surgery, and severe postoperative complication). The last binary feature, i.e. severe postoperative complication, was used as output variable for the supervised ML classifiers. All the continuous features were standardized to have mean 0 and standard deviation 1 before they enter the classifier. HbA1c was log transformed before standardization because of its asymmetric distribution.

### Descriptive and inferential statistical methods

Demographic and baseline characteristics of the patients were presented using descriptive statistical methods. Continuous variables were portrayed as mean and standard deviation (SD), or median and interquartile range where suitable, while categorical variables were outlined as counts and percentages. The difference between the patient presenting and without severe postoperative complication was tested using the Student’s t-test or the Mann-Whitney U test for normally or asymmetrically distributed continuous variables, respectively; and χ^2^ test was used for binary variables.

### ML algorithms

In current study, eight base ML algorithms, i.e. logistic regression, linear discriminant analysis (LDA), quadratic discriminant analysis (QDA), decision tree, k-nearest neighbor (KNN), support vector machine (SVM), multilayer perceptron (MLP) and deep learning neural network (NN), and 11 ensemble algorithms, i.e. adaptive boosting (AdaBoost) logistic regression, bagging LDA, bagging QDA, random forest, extremely randomized (Extra) trees, AdaBoost Extra trees, gradient regression tree, AdaBoost Gradient trees, bagging KNN, AdaBoost SVM, and bagging MLP, were implemented.

### Ensemble learning

In order to improve generalizability and robustness over a single ML algorithm, we also used ensemble methods to combine multiple base or ensemble algorithms. Five ensemble methods were applied in our study:

- AdaBoost for logistic regression, Extra trees, gradient regression trees, and SVM(Schapire, 2003);
- bagging for LDA, QDA, KNN, and MLP(Kotsiantis et al., 2007);
- random forests for decision tree(Liaw & Wiener, 2002);
- Extra trees for decision tree(Geurts et al., 2006);
- gradient boosted regression trees for decision tree(Friedman, 2002).

### Initialization and optimization of hyperparameters

ML algorithms involve a number of hyperparameters that have to be fixed before running the algorithms. In contrast to the parameters that are learned by training, hyperparameters determine the structure of a ML algorithm and how the algorithm is trained. The initial values of the hyperparmeters for each ML algorithm used in our study are the default values specified in the employed software packages based on recommendations or experience(Probst et al., 2018). In KNN algorithm, ten nearest neighbors were used. In MLP algorithm, two hidden layer were used with five and two neurons, respectively. In deep learning NN algorithm, the sequential linear stack of layers was used, with five hidden layers (three dense layers and two dropout regularization layers). For the detailed hyperparameterization of the algorithms, please refer the scikit-learn user manual at http://scikit-learn.org/stable/supervised_learning.html(Pedregosa et al., 2011) and the Keras Documentation at https://keras.io/.

The hyperparameter optimization is defined as a tuple of hyperparameters that yields an optimal algorithm which minimizes a predefined loss function (i.e. cross entropy loss function in our study, see Annex 1) on a held-out validation set of the training data. The most wildly used however exhaustive grid search was used to perform hyperparameter optimization in our study, which specified subset of the hyperparameter space of a ML algorithm and was evaluated by cross-validation using the training data(Bergstra & Bengio, 2012).

### Cross validation

For training data, *k*-fold (*k* = 5 in our analyses) cross-validated predictions were used as predicted values. This approach involves randomly dividing the training data into *k* groups, or folds, of approximately equal size. Then an algorithm is trained on the *k*-1 folds and the rest one fold is retained as the validation fold for testing the algorithm. The process is repeated until the algorithm is validated on all the *k* folds. For each patient in the training data, the predicted value that he/she obtained is the prediction when he/she was in the validation fold. Therefore, only cross-validation strategies that assign all patients to a validation fold exactly once can be used for the cross-validated prediction(James et al., 2013).

## SMOTE

The bariatric surgery data is extreme imbalanced, i.e. only 1,408 of 44,061 (3.2%) patients experienced severe postoperative complication after bariatric surgery. The imbalance often results in serious bias in the performance metrics(Batista et al., 2004). Therefore, we performed synthetic minority oversampling technique (SMOTE) to tackle the imbalance(Chawla et al., 2002). SMOTE generates a synthetic instance by interpolating *m* instances (for a given integer value *m*) of the minority class that lies close enough to each other to achieve the desired ratio between the majority and minority classes. In our study, a 1:1 ratio between the patients presenting severe postoperative complication and without severe postoperative complication was achieved in the training data, i.e. SMOTE training data. The aforementioned nine of the 11 ensemble ML algorithms and the deep learning NN were also implemented for the SMOTE training data.

### Performance metrics

The performance of the in total 29 ML algorithms were evaluated using accuracy, sensitivity, specificity, and area under the receiver operating characteristic (ROC) curve. ROC curve shows the trade-off that the algorithms set the different threshold values for the posterior probability for the prediction.

Terminology and derivations of accuracy, sensitivity, specificity, and area under the ROC curve are given in Annex 1.

### Software and hardware

The descriptive and inferential statistical analyses were performed using Stata 15.1 (StataCorp, College Station). ML algorithms were achieved using packages scikit-learn 0.19.1 (scikit-learn, http://scikit-learn.org/)(Pedregosa et al., 2011) and Keras 2.1.6 (Keras, https://keras.io/) in Python 3.6 (Python Software Foundation, https://www.python.org/).

All the computation was conducted in a computer with 64-bit Windows 7 Enterprise operation system (Service Pack 1), Intel ^®^ Core ^TM^ i5-4210U CPU @ 2.40 GHz, and 16.0 GB installed random access memory.

## Author contributions

Yang Cao, Xin Fang, Conceptualization, Data curation, Software, Formal analysis, Writing-original draft; Yang Cao, Supervision, Investigation, Visualization, Methodology; Erik Stenberg Data curation, Conceptualization, Validation, Investigation, Writing-original draft; Johan Ottosson, Erik Näslund, Investigation, Writing-original draft.

## Ethics

The study was approved by the Regional Ethics Committee in Stockholm and was conducted in accordance with the ethical standards of the Helsinki Declaration (6th revision).

**Table 1.** Base line characteristics of the training patients

|  | All<br>N=37,811 | No serious<br>complication<br>N=36,591<br>(96.8%) | Having serious<br>complication<br>N=1,220 (3.2%) | p-value |
| --- | --- | --- | --- | --- |
| Age in years, mean ± SD | 41.2 ± 11.2 | 41.1 ± 11.2 | 42.9 ± 10.7 | <0.001* |
| Sex, n (%) |  |  |  |  |
| Female | 28,682 (75.9%) | 27,766 (75.9%) | 916 (75.1%) | 0.521<sup>†</sup> |
| Male | 9,129 (24.1%) | 8,825 (24.1%) | 304 (24.9%) |  |
| BMI in kg/m<sup>2</sup>, mean ± SD | 42.12 ± 5.66 | 42.13 ± 5.66 | 41.79 ± 5.58 | 0.0355* |
| WC in cm, mean ± SD | 126.0 ± 14.0 | 126.0 ± 14.0 | 126.2 ± 13.8 | 0.6018* |
| HbA1c, median (P25, P75) | 38 (35, 42) | 38 (38, 32) | 38 (35, 43) | 0.0090<sup>‡</sup> |
| Comorbidity, n (%) |  |  |  |  |
| Sleep apnoea | 3,792 (10.0%) | 3,656 (10.0%) | 136 (11.2%) | 0.186<sup>†</sup> |
| Hypertension | 9,760 (25.8%) | 9,404 (25.7%) | 356 (29.2%) | 0.006<sup>†</sup> |
| Diabetes | 5,407 (14.3%) | 5,204 (14.2%) | 203 (16.6%) | 0.018<sup>†</sup> |
| Dyslipidaemia | 3,802 (10.1%) | 3,667 (10.0%) | 135(11.1%) | 0.233<sup>†</sup> |
| Dyspepsia | 3,970 (10.5%) | 3,803 (10.4%) | 167 (13.7%) | <0.001<sup>†</sup> |
| Depression | 5,609 (14.8%) | 5,409 (14.8%) | 200 (16.4%) | 0.119<sup>†</sup> |
| Musculoskeletal pain | 4,905 (13.0%) | 4,754 (13.0%) | 151 (12.4%) | 0.529<sup>†</sup> |
| Previous venous thromboembolism | 918 (2.4%) | 875 (2.39%) | 43 (3.52%) | 0.011<sup>†</sup> |
| Revisional surgery | 1,367 (3.6%) | 1,261 (3.5%) | 106 (8.7%) | <0.001<sup>†</sup> |

SD: standard deviation; BMI, body mass index; WC, waist circumference; P25, the 25th percentile;
P75, the 75th percentile.
\*t-test was used; <sup>†</sup> $\chi^2$ test was used; <sup>‡</sup>Mann-Whitney U test was used.

**Table 2.** Base line characteristics of the test patients

|  | All<br>N=6,250 | No serious<br>complication<br>N=6,062 (97.0%) | Having serious<br>complication<br>N=188 (3.0%) | p-value |
| --- | --- | --- | --- | --- |
| Age in years, mean ± SD | 41.2 ± 11.5 | 41.2 ± 11.5 | 42.9 ± 11.8 | 0.0423<sup>*</sup> |
| Sex, n (%) |  |  |  |  |
| Female | 4,832 (77.3%) | 4,682 (77.2%) | 150 (79.8%) | 0.411<sup>†</sup> |
| Male | 1,418 (22.7%) | 1,380 (22.8%) | 38 (20.2%) |  |
| BMI in kg/m<sup>2</sup>, mean ± SD | 41.22 ± 5.87 | 41.20 ± 5.89 | 41.95 ± 5.40 | 0.0848<sup>*</sup> |
| WC in cm, mean ± SD | 123.3 ± 14.1 | 123.2 ± 14.0 | 126.2 ± 14.7 | 0.0086<sup>*</sup> |
| HbA1c, median (P25,<br>P75) | 37 (34, 41) | 37 (34, 41) | 38 (35, 44) | 0.0017<sup>‡</sup> |
| Comorbidity, n (%) |  |  |  |  |
| Sleep apnoea | 622 (10.0%) | 607 (10.0%) | 15 (8.0%) | 0.359<sup>†</sup> |
| Hypertension | 1,563 (25.0%) | 1,506 (24.8%) | 57 (30.3%) | 0.088<sup>†</sup> |
| Diabetes | 761 (12.2%) | 734 (12.1%) | 27 (14.4%) | 0.352<sup>†</sup> |
| Dyslipidaemia | 518 (8.3%) | 493 (8.13%) | 25 (13.3%) | 0.011<sup>†</sup> |
| Dyspepsia | 645 (10.3%) | 620 (10.2%) | 25 (13.3%) | 0.173<sup>†</sup> |
| Depression | 1,096 (17.5%) | 1,053 (17.4%) | 43 (22.9%) | 0.051<sup>†</sup> |
| Musculoskeletal pain | 1,315 (21.0%) | 1,268 (20.9%) | 47 (25.0%) | 0.176<sup>†</sup> |
| Previous venous<br>thromboembolism | 182 (2.9%) | 177 (2.99%) | 5 (2.7%) | 0.834<sup>†</sup> |
| Revisional surgery | 61 (1.0%) | 54 (0.9%) | 7 (3.7%) | <0.001<sup>†</sup> |

SD: standard deviation; BMI, body mass index; WC, waist circumference; P25, the 25th percentile;
P75, the 75th percentile.
<sup>\*</sup>t-test was used; <sup>†</sup> $\chi^2$ test was used; <sup>‡</sup>Mann-Whitney U test was used.

**Table 3.** Performance of the algorithms

| Algorithm | Training data |  |  | Test data |  |  |
| --- | --- | --- | --- | --- | --- | --- |
|  | Accuracy (%) | Specificity | Sensitivity | Accuracy (%) | Specificity | Sensitivity |
| Logistic | 96.9 | 1.000 | 0.000 | 97.1 | 1.000 | 0.000 |
| AdaBoost Logistic | 96.9 | 1.000 | 0.000 | 97.1 | 1.000 | 0.000 |
| Oversampling AdaBoost Logistic | 46.5 | 0.382 | 0.547 | 76.9 | 0.786 | 0.227 |
| LDA | 96.9 | 1.000 | 0.000 | 97.1 | 1.000 | 0.000 |
| Bagging LDA | 96.9 | 1.000 | 0.000 | 97.1 | 1.000 | 0.000 |
| Oversampling Bagging LDA | 46.3 | 0.370 | 0.556 | 79.0 | 0.807 | 0.212 |
| QDA | 92.8 | 0.954 | 0.107 | 94.7 | 0.973 | 0.076 |
| Bagging QDA | 93.2 | 0.958 | 0.103 | 95.5 | 0.982 | 0.068 |
| Oversampling bagging QDA | 55.4 | 0.401 | 0.707 | 56.1 | 0.566 | 0.417 |
| Decision tree | 93.5 | 0.963 | 0.038 | 93.1 | 0.958 | 0.045 |
| Random Forest | 96.9 | 1.000 | 0.000 | 97.0 | 1.000 | 0.000 |
| Oversampling Random Forest | 94.5 | 0.925 | 0.965 | 96.6 | 0.995 | 0.008 |
| ExtRa Trees | 96.6 | 0.997 | 0.006 | 96.7 | 0.996 | 0.015 |
| AdaBoost ExtRa Trees | 96.6 | 0.996 | 0.004 | 96.6 | 0.995 | 0.008 |
| Oversampling AdaExtra Trees | 93.0 | 0.881 | 0.980 | 95.3 | 0.982 | 0.015 |
| Gradient regression trees | 96.9 | 1.000 | 0.000 | 97.1 | 1.000 | 0.008 |
| AdaBoost Gradient trees | 96.8 | 0.998 | 0.000 | 97.0 | 0.999 | 0.000 |
| Oversampling AdaGradient trees | 97.0 | 0.972 | 0.968 | 97.0 | 0.999 | 0.000 |
| KNN | 96.9 | 1.000 | 0.000 | 97.1 | 1.000 | 0.000 |
| Bagging KNN | 96.9 | 1.000 | 0.000 | 97.1 | 1.000 | 0.000 |
| Oversampling Bagging KNN | 79.4 | 0.592 | 0.996 | 82.3 | 0.841 | 0.235 |
| SVM | 96.9 | 1.000 | 0.000 | 97.1 | 1.000 | 0.000 |
| AdaBoost SVM | 96.9 | 1.000 | 0.000 | 97.1 | 1.000 | 0.000 |
| Oversampling AdaBoost SVM | 53.6 | 0.397 | 0.675 | 60.6 | 0.614 | 0.364 |
| MLP | 96.9 | 1.000 | 0.000 | 97.1 | 1.000 | 0.000 |
| Bagging MLP | 96.9 | 1.000 | 0.000 | 97.1 | 1.000 | 0.000 |
| Oversampling bagging MLP | 45.7 | 0.226 | 0.687 | 96.6 | 0.994 | 0.015 |
| Deep learning NN | 96.9 | 1.000 | 0.000 | 97.1 | 1.000 | 0.000 |
| Oversampling deep learning NN | 62.1 | 0.484 | 0.757 | 93.3 | 0.959 | 0.068 |

AdaBoost, adaptive boosting; LDA, linear discriminant analysis; QDA, quadratic discriminant analysis; ExtRa, extremely randomized; AdaExtra, adaptive boosting extremely randomized; AdaGradient, adaptive boosting gradient; KNN, k-nearest neighbor; SVM, support vector machine; MLP, multilayer perceptron; NN, neural network

**Figure 1.**
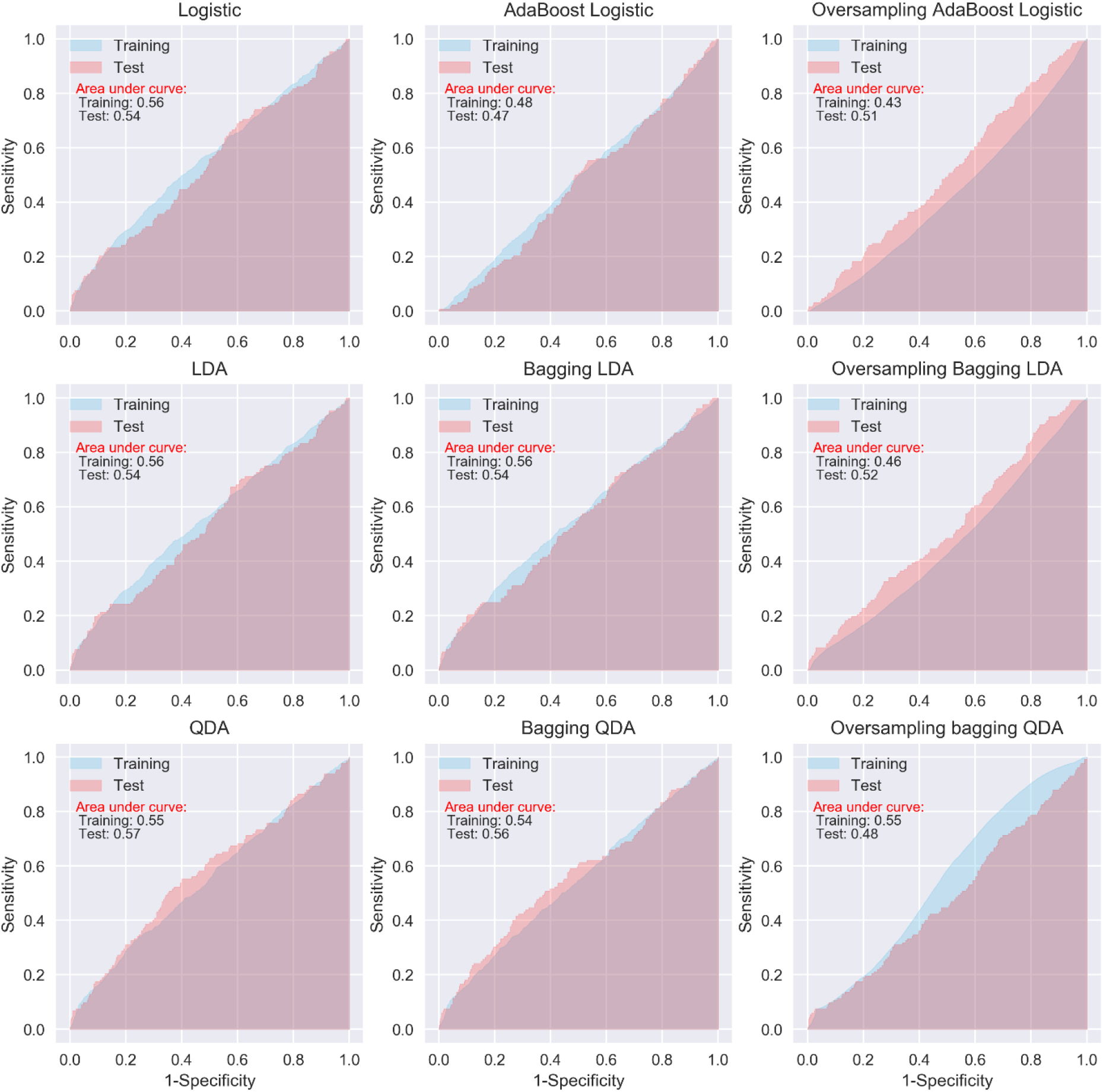
ROC curves of logistic regression, LDA and QDA

**Figure 2.**
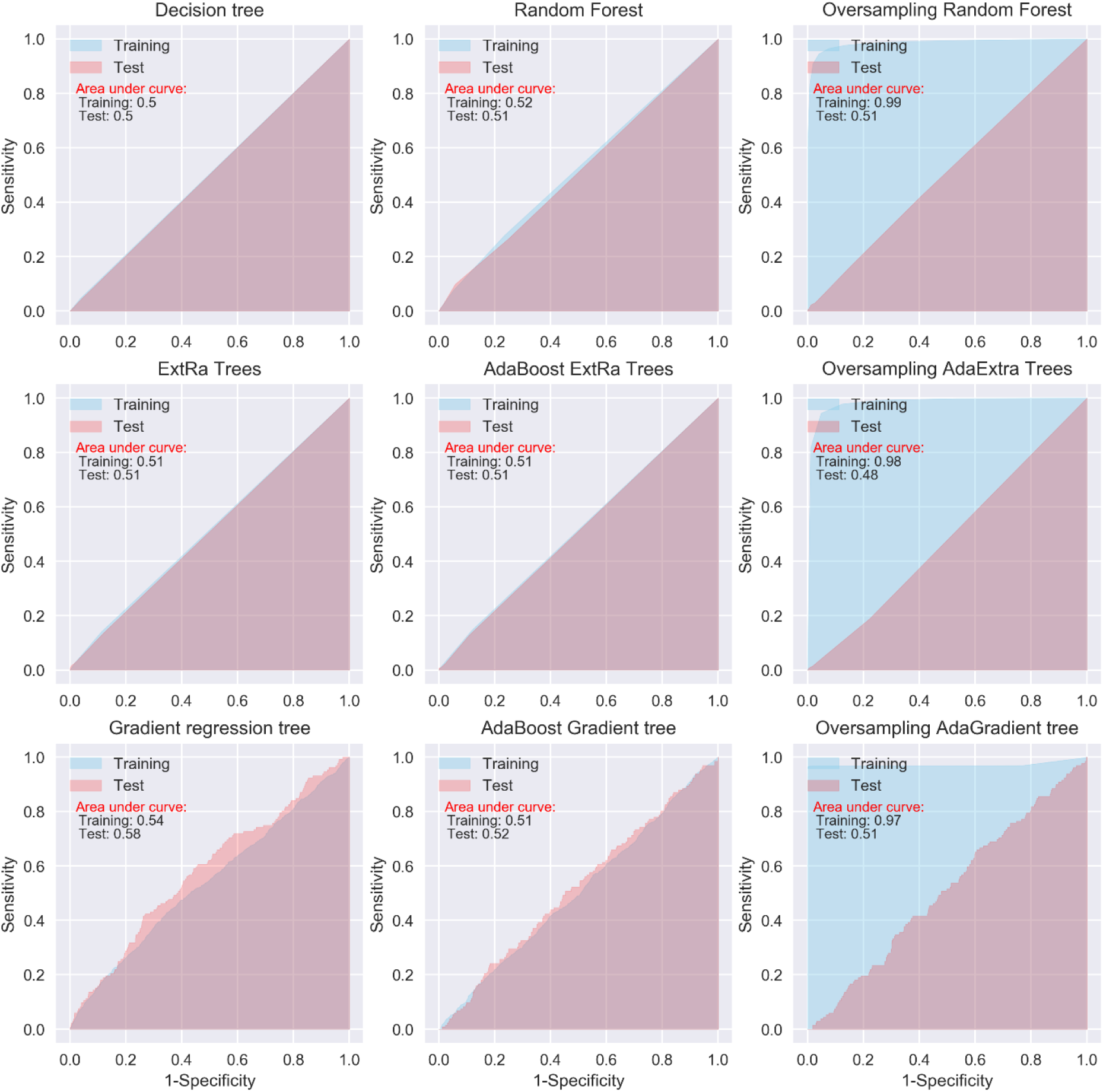
ROC curves of tree-based algorithms

**Figure 3.**
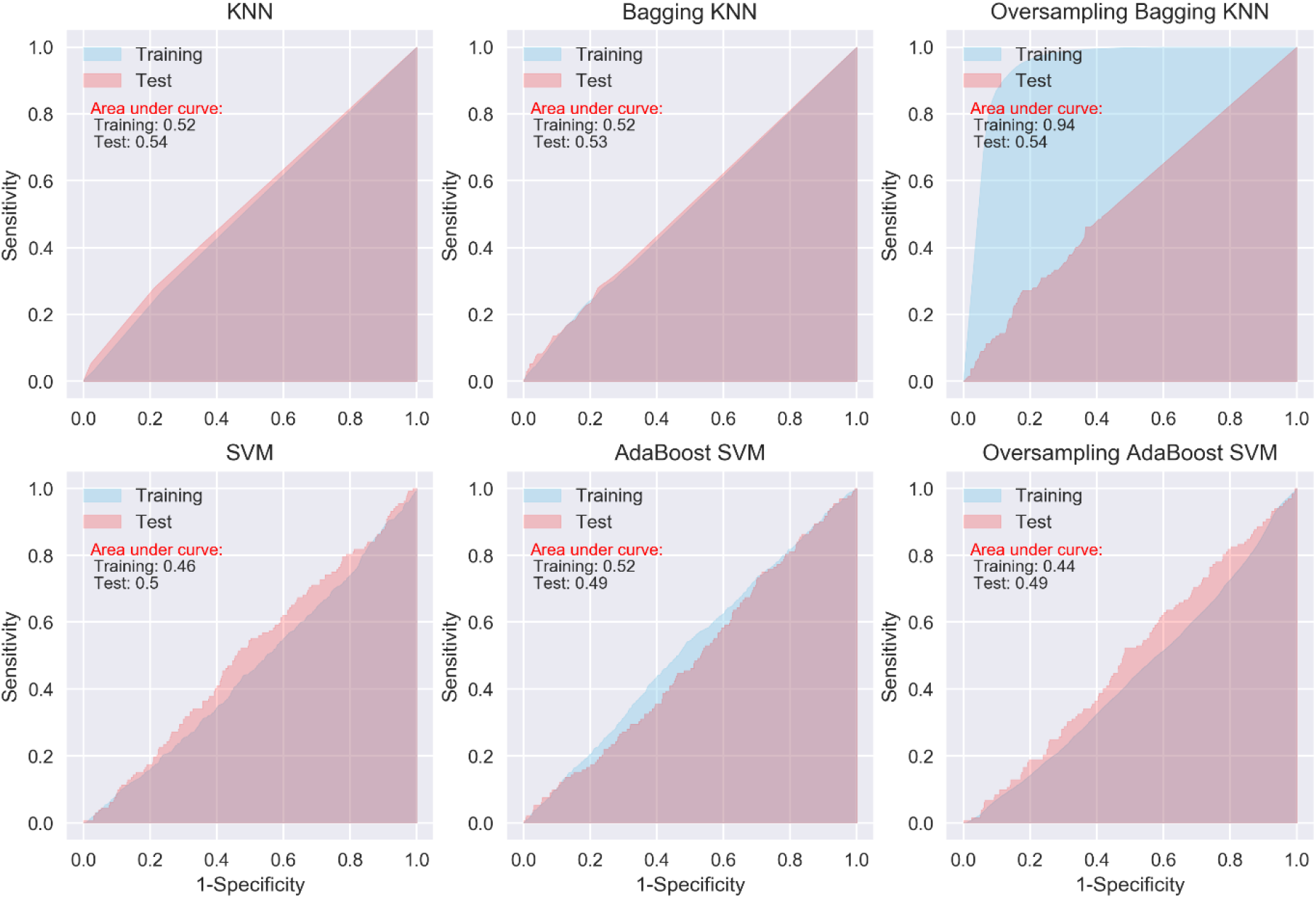
ROC curves of KNN and SVM

**Figure 4.**
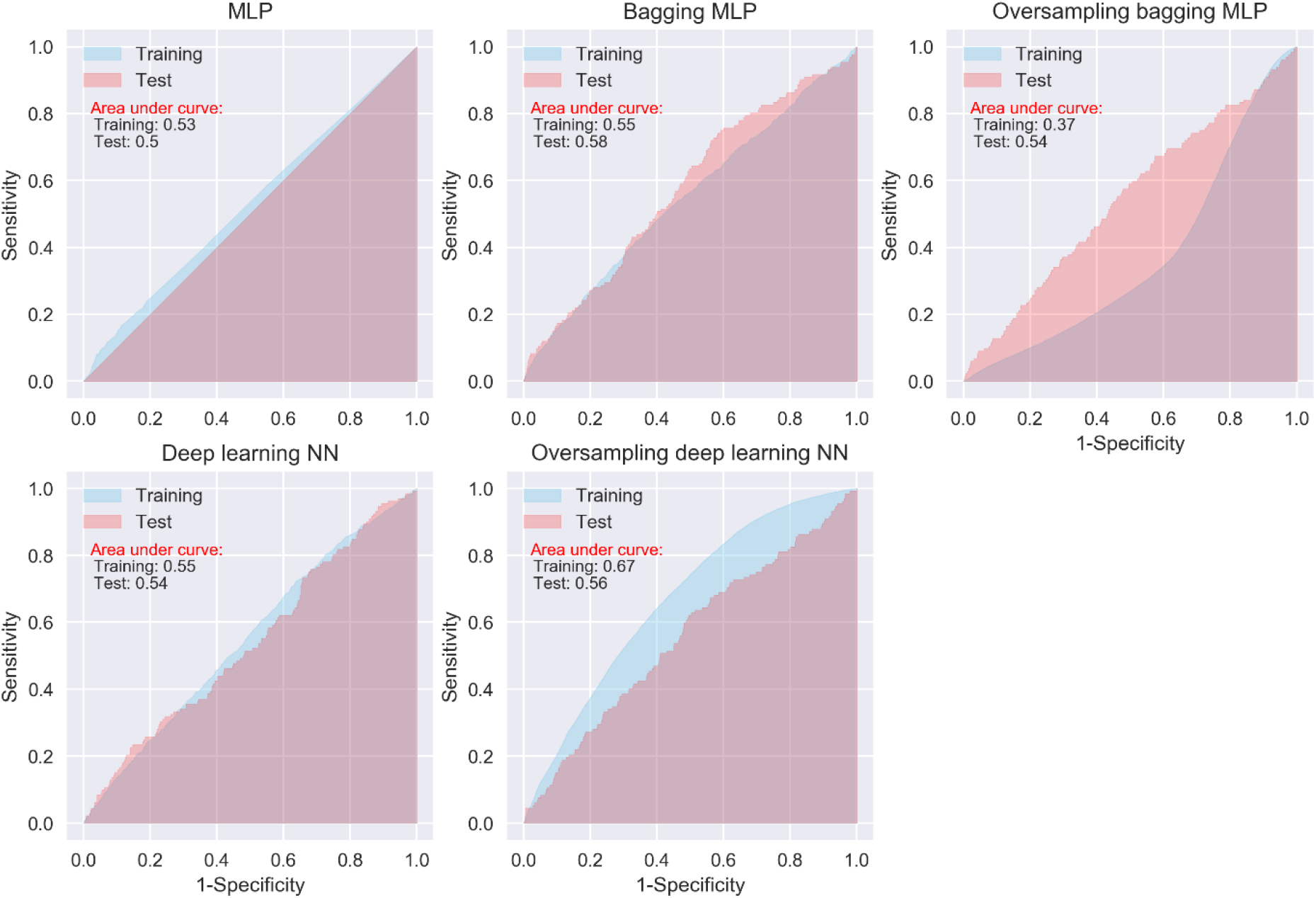
ROC curves of neural network algorithms

## Annex 1: Terminology and derivations

Cross Entropy loss (Log loss):

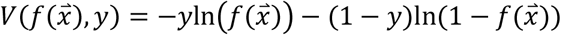

where *y* is true classifier ∈ {0, 1} and 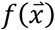 is predicted value.

True or false refers to the predicted outcome being correct or wrong, while positive or negative refers to presenting severe complication or no severe complication.

ACC: accuracy
SEN: sensitivity
SPE: specificity
TP: number of true positives, i.e. patient presenting severe complication correctly predicted as positive
TN: number of true negatives, i.e. patient without severe complication correctly predicted as negative
FP: number of false positives, i.e. patient without severe complication wrongly predicted as positive
FN: number of false negatives, i.e. patient presenting severe complication wrongly predicted as negative
Total: total number of the patients, i.e. TP+TN+FP+FN
P: number of patients presenting severe complication, i.e. TP+FN
N: number of patients without severs complication, i.e. TN+FP
AUC: area under the receiver operating characteristic (ROC) curve for binary outcome
T: threshold for a patient is classified as presenting severe complication if X>T, where X is predicted probability of a patients presenting severe complication by an algorithm.

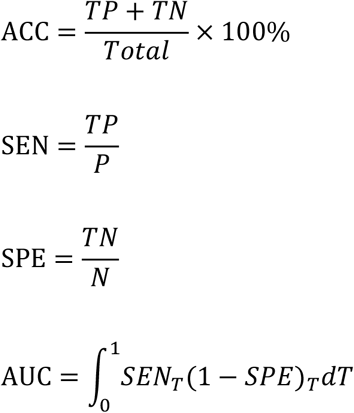

